# Tumor Organoids Modeling Reveals Timed Responses and Interplay of Radiotherapy and Chemotherapy in Pancreatic Cancer

**DOI:** 10.1101/2025.09.21.677544

**Authors:** Chayu Yang, Zachery Keepers, Hem D Shukla, Lei Ren

## Abstract

**Purpose:** Model the therapeutic effects of chemotherapy, radiotherapy, and chemoradiotherapy to study the heterogeneous responses of patient-derived tumor organoids (PDTOs).

**Methods:** We proposed novel mathematical forecasting frameworks based on logistic growth ordinary differential equations (ODEs) and the characteristics of tumor responses to model the therapeutic effects of chemo- and radiation-induced killing. To validate the models, we cultured PDTOs from different patients and treated them with radiation (4 Gy and 8 Gy), chemotherapy (FOLFIRINOX at a previously determined organoid-specific IC50 dose), and combined regimens, respectively. The diameters of 20-40 organoids per patient were tracked and measured using brightfield images up to 7 or 9 days following each treatment to capture organoid growth dynamics data, which was used for model fitting. The accuracy of the modeling was evaluated by the average normalized mean squared errors (NMSE) of data fitting.

**Results:** The proposed mathematical modeling frameworks accurately captured the observed growth dynamics of three PDTO samples after chemotherapy, radiotherapy, and chemoradiotherapy, as reflected by the average NMSE of data fitting, which are all close to zero (less than 0.0045). The fitted parameters, including killing strength, effect window, and peak killing timing, revealed significant heterogeneities in treatment responses across different PDTOs. Chemotherapy shows pronounced effectiveness in the early stage, while radiotherapy exhibits the effect later in the first week, and a significantly stronger secondary response occurs one week after radiation. Chemoradiotherapy combines the strengths of both modalities, producing pronounced effects in both response phases, with modeling results suggesting that the radiation-induced killing effect may play a dominant role in the synergistic interaction.

**Conclusions:** Our modeling frameworks demonstrated high accuracy in modeling the heterogeneous therapeutic responses of PDTOs and provided insights into the dynamic killing effects and interplay between chemotherapy and radiotherapy. The modeling of therapeutic responses of PDTOs provides a valuable tool for optimizing treatment regimens and informing clinical trial design, thereby improving the efficacy of personalized medicine.

## 1. Introduction

In recent years, patient-derived tumor organoids (PDTOs) have emerged as transformative in vitro models in cancer research, revolutionizing the landscape of preclinical oncology. Driven by advances in stem cell and three-dimensional (3D) culture technologies, organoids enable the rapid generation of 3D tumor tissue that replicates the tumor microenvironment and key biological features of the original patient tumor, including genetic, histological, and phenotypic heterogeneity [1, 2, 3]. Unlike conventional 2D cell lines or patient-derived xenografts, organoids can provide superior physiological relevance and reproducibility in a matter of weeks following tumor biopsy or resection [3, 4]. PDTOs have now been successfully developed from a wide spectrum of human cancers, including but not limited to colorectal, pancreatic, breast, lung, liver, and prostate tumors. These models are particularly valuable for capturing patient-specific tumor behavior, including inter- and intra-tumoral heterogeneity, and for assessing responses to chemotherapy, radiotherapy, and targeted therapies [5, 6, 7]. As such, they can serve as critical platforms for in vitro drug screening, biomarker identification, and therapeutic optimization. Importantly, PDTOs are increasingly integrated into translational and personalized medicine pipelines, offering a powerful means to evaluate mutation-specific treatment strategies, simulate tumor evolution under therapeutic pressure, and reduce reliance on expensive and time-consuming clinical trials. Continued improvements in organoid culture protocols, the use of tissue-specific growth factors, and the expansion of biobanking capacity have further accelerated their application across research and clinical settings [7, 8, 9].

An expanding body of research underscores the value of PDTOs in modeling therapeutic responses to chemotherapy, radiotherapy, and chemoradiotherapy (CRT). Tumor organoids derived from individual patients have consistently demonstrated treatment sensitivity or resistance patterns that align with observed clinical outcomes, reinforcing their potential as ex vivo predictive tools for personalized therapy design [10-12]. Radiotherapy studies using PDTOs from rectal, pancreatic, and head and neck cancers have revealed complex temporal dynamics, including progressive tumor shrinkage, delayed DNA damage repair, and variable recovery potential over time [13]. Similarly, chemotherapy screening using agents such as 5-fluorouracil, oxaliplatin, and gemcitabine has provided valuable insights into patient-specific dose-response relationships and intra-tumoral heterogeneity [14]. When exposed to combined treatment modalities, such as sequential or concurrent chemoradiotherapy, PDTOs frequently exhibit nonlinear response behaviors and synergistic effects that are not easily discernible through experimental methods alone. These complex interactions underscore the importance of integrating mathematical modeling to decode response dynamics and optimize treatment regimens [15, 16].

Mathematical modeling has a rich history in oncology, serving as a quantitative tool to provide insights into tumor growth kinetics, treatment response, and evolutionary dynamics. Early growth formulations, such as the logistic and Gompertz models, were widely used to describe organismal and in vivo tumor volume trajectories, often showing superior predictive power in forecasting future tumor behavior and regression patterns [17]. Classic ODE-based growth models can be extended naturally to organoid systems, and more sophisticated variants incorporate treatment-induced effects, such as cell death, damage repair, cell-cycle arrest, decay, or clearance of apoptotic and necrotic debris, spatial limitations, and nutrient diffusion. These enhancements aim to bridge the gap between in vitro observations and clinical behavior, enabling richer interpretations of organoid data. However, despite these advancements, the mathematical modeling of treatment-induced effects in the organoid system remains largely underexplored.

In this study, we developed mathematical models based on logistic growth to describe the therapeutic effect, which is represented by the rate of cell killing. To acquire data for model fitting, we cultured the tumor organoids from three pancreatic cancer patients and treated them with radiotherapy, chemotherapy, and combined chemoradiotherapy, respectively. The dynamic changes of organoid sizes after treatment were then measured to fit and examine the proposed models. We aim to integrate mathematical modeling with data from cultured PDTOs to estimate hidden biological parameters, predict future tumor behavior under various treatment scenarios, quantify dynamic and synergistic treatment effects not directly observable in experiments, and support personalized treatment planning by predicting patient responses based on PDTO modeling. This integrative approach helps bridge preclinical testing with more effective and individualized therapies in the clinic, providing a much-needed tool to optimize treatment strategies to improve patient outcomes.

## 2. Methods and Materials

### 2.1 Pancreatic Tumor Organoid Culture and Subculture

Patient-derived pancreatic tumor organoids were obtained from the National Cancer Institute’s Patient-Derived Models Repository (PDMR). All media components were as detailed in SOP30101 (NCI, 2023). Upon receipt, cryovials containing organoids were warmed in a water bath at 37 °C for 1-2 minutes, washed with wash media (NCI, 2023) and centrifuged at 300g for 5 minutes twice, and resuspended in Cultrex™ PathClear Reduced Growth Factor BME, Type 2 (R&D Systems, #3533-005-02). Organoids were then plated onto 24-well tissue culture plates (Costar, #3524) that had been pre-warmed for more than 24 hours at 37 °C. Tissue culture plates were flipped and incubated at 37 °C for 15 minutes to allow three-dimensional distribution of organoids throughout BME as it polymerized in the shape of a dome. After incubation, 750 μL of panc complete (NCI, 2023) media was added to each well. Organoids were propagated and subcultured three times to generate stock, which was stored in liquid nitrogen at −180 °C. All experiments utilized organoids from the original stock and were passaged twice from cryopreservation before receiving any experimental exposure. To passage organoids, a volume of splitting media (NCI, 2023) equal to the volume of media in the culture well was added, followed by incubation at 37 °C for 60-90 minutes. Domes were disrupted with gentle pipetting and organoids were transferred for washing, centrifugation, and subculture as described above.

### 2.2 Treatment with Ionizing Radiation, Chemotherapy Drugs, and Combination Regimens

Tumor organoids obtained from three different patients, labeled #7800, #8510, and #11777, were dissociated into single cells using Accutase (Innovative Cell Technologies, #AT-104) and plated as in 30 μL BME2 domes with 2500 cells per dome. Treatment groups included ionizing radiation at 4 Gy and 8 Gy, FOLFIRINOX was administered at a previously determined organoid-specific IC50 dose, and combinations of radiation and FOLFIRINOX. The four components of FOLFIRINOX were mixed as a previously determined ratio (n): 4 μM of 5FU (Sigma-Aldrich, #F6627), 12.5 nM of SN38 (ChemSelleck, #S4908), 0.5 μM of oxaliplatin (ChemSelleck, #S1224), and 2 μM of leucovorin (Sigma-Aldrich, #PHR1541). Brightfield images were acquired using an EVOS XI light microscope (ThermoFisher Scientific, USA) starting on day 0 post-treatment for all three organoid samples and lasting to day 7 or day 9. For each sample in each group, 20–40 organoids of sufficient size were traced from the images, and their sizes were measured using MATLAB image processing tools to characterize the growth dynamics.

### 2.3 Mathematical modeling

Due to competition for limited nutrients and environmental constraints in organoid cultures, we adopt the logistic growth model [17] to characterize the cell proliferation of an organoid. The model describes a self-limiting growth of a biological population with an intrinsic growth rate *λ* that decreases linearly to zero as the population approaches carrying capacity *K* over time *t*. The logistic growth model is given by the following ordinary differential equation

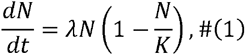

where *N* is the total population size at time t. For chemotherapy and radiotherapy, we introduce functions C(t) and R(t) to represent the time-dependent chemo-induced and radiation-induced killing rates, respectively. Hence the growth model with chemotherapy can be given by

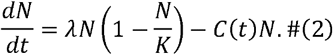

Following radiation exposure, the cell population can be divided into a surviving fraction and a non-surviving fraction, and we denote these two fractions as active cells (*A*) and inactive cells (*I*), respectively. The initial cell populations of these two fractions can be estimated using the linear-quadratic model

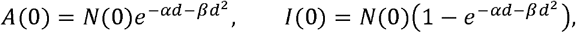

where *α* and *β* are radiosensitivity parameters and *d* is the radiation dose. In the radiotherapy growth model, we assume the inactive (non-surviving) cells do not proliferate and are cleared at an average constant rate *μ*, primarily through apoptosis, necrosis, and related processes. The model is thus given by

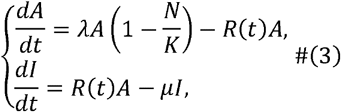

where *A* and *I* represent the number of active cells and inactive cells, respectively, and parameters *λ* and *μ* defines the growth rate of active cells and the removal rate of inactive cells, respectively. Based on the growth models (2) and (3), the cell population growth dynamics after a combination of radiotherapy and chemotherapy can be modeled as

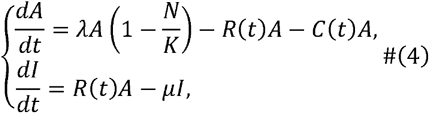

where we assume that the chemo-induced killing effect primarily acts on active cells and has no contribution to inactive cells.

Given that drug exposure induces immediate cytotoxic and/or cytostatic effects during the first few days, followed by a plateau phase and potential regrowth [11, 18], we assume that the chemo-induced killing effect initially increases to a maximum before gradually declining to zero. Accordingly, we model the chemo-induced killing rate as

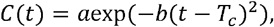

where *a, b*, and *T*_*c*_ are parameters representing drug sensitivity, the duration of killing effect window, and the timing of the peak killing effect, respectively. In this framework, a larger *a* corresponds to a stronger drug sensitivity, a larger *b* indicates a shorter effective killing window, and a larger *T*_*c*_ reflects a delayed peak response to treatment. In contrast to chemotherapy, organoids typically exhibit limited early-phase sensitivity to radiotherapy. A clear response generally emerges within the first week, with cell killing potentially persisting and intensifying beyond this period [19, 20]. To capture the strengths and timings of these responses, we assume that two distinct waves of radiation-induced killing occur at time points *T*_1_ and *T*_2_, with respective maximum strengths *u*_1_and *u*_2_. We then set the radiation-induced killing rate to be

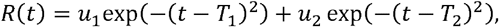

where *u*_1_ and *u*_2_ represent parameters related to radiosensitivity, and *T*_1_ and *T*_2_ specify the timings of the two local maxima in the radiation-induced killing effect.

### 2.4 Model data fitting

Since the mathematical models we developed are intended to simulate the temporal dynamics of cell numbers of an organoid, the collected diameter measurements must first be converted into cell number estimates before fitting the models to the dataset. Assuming each organoid is spherical, the number of cells of an organoid with diameter *D* can be approximated by 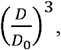 Where *D*_0_ is the average cell diameter, estimated as 22.5 *μm* for pancreatic tumor cells [21], and *D* is calculated from the organoid area that was measured using MATLAB image processing, as shown in **Figure S1** of the Supplementary Material. From this, the maximum diameter *D_K_* of an organoid when it reaches the carrying capacity *K* is given by *D*_*K*_ = *D*_0_*K*^1/3^. Model fitting for each organoid is performed using the nonlinear least-squares solver *lsqcurvefit* in MATLAB, which estimates parameter values by minimizing the sum of squared errors 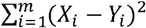 between the model predictions (*Y*_1_, *Y*_2_ … *Y*_*m*_) and the collected dataset (*X*_1_, *X*_2_ … *X*_*m*_) at time points (*t*_1_, *t*_2_ … *t*_*m*_) where *X*_*i*_ and *Y*_*i*_ represent the estimated and predicted cell numbers at time *t*_*i*_ respectively, and m denotes the total number of data points for an organoid. To account for size variations among organoids, we use the normalized mean squared error 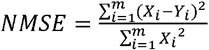, an estimate of the overall deviation between the predicted and measured values, to assess the goodness of fit for each organoid. The NMSE provides a standardized measure of how accurately the model captures the observed dynamics. Generally, a lower value of NMSE indicates a better-fitting result.

## 3. Results

### 3.1 Control group

We first applied model (1) to fit the control groups of PDTOs without treatments, thereby estimating their overall growth rates and average maximum diameters in the cultured environment. **Figure 1(A)** shows the fitting curves alongside the collected data for organoids #7800, while the corresponding results for #8510 and #11777 are provided in **Figures S2(A)** and **S3(A)**. The parameter estimates, together with their 95% confidence intervals (CIs) and measures of goodness of fit, are summarized in **Table 1 (Control)**. The fits are of similar quality, with an average NMSE of approximately 0.0030, and the poorest case exhibits a maximum NMSE of only 0.0095 among all control-group organoids. Among the three samples, organoids #11777 show the lowest average growth rate (0.431 per day) and an average maximum diameter of 360.8 *μm*, whereas organoids #8510 display the fastest average growth rate (0.657 per day) and the largest average maximum diameter (406.5 *μm*). Using the 99.9% CI of the maximum diameter to estimate the largest size achievable by any organoid within a sample, the maximum sizes for the organoids #7800, #8510, and #11777 are approximately 578.8 *μm*, 700.3 *μm*, and 695.8 *μm*, respectively.

**Table 1.** Fitting parameter values with 95% CIs and goodness of fit in the control, chemotherapy, and radiotherapy groups.

|  | PDTOs ID | #7800 | #8510 | #11777 |
| --- | --- | --- | --- | --- |
| Control | $\lambda$ (day<sup>-1</sup>) | $0.464 \pm 0.033$ | $0.657 \pm 0.053$ | $0.431 \pm 0.047$ |
| | $D_K$ ($\mu m$) | $302.8 \pm 155.2$ | $406.5 \pm 164.2$ | $360.8 \pm 187.3$ |
|  | Ave. NMSE | 0.0028 | 0.0043 | 0.0017 |
|  | Max. NMSE | 0.0078 | 0.0095 | 0.0094 |
| Chemotherapy | $a$ (day<sup>-1</sup>) | $0.515 \pm 0.037$ | $0.906 \pm 0.122$ | $0.395 \pm 0.026$ |
| | $b$ | $0.077 \pm 0.019$ | $0.162 \pm 0.085$ | $0.072 \pm 0.028$ |
| | $T_c$ (day) | $3.5 \pm 0.3$ | $4.2 \pm 0.3$ | $5.3 \pm 1.0$ |
|  | Ave. NMSE | 0.0010 | 0.0023 | 0.0013 |
|  | Max. NMSE | 0.0038 | 0.0062 | 0.0068 |
| Radiotherapy<br>at 4 Gy | $\lambda$ (day<sup>-1</sup>) | $0.397 \pm 0.033$ | $0.773 \pm 0.076$ | $0.412 \pm 0.028$ |
| | $u_1$ (day<sup>-1</sup>) | $0.475 \pm 0.090$ | $1.636 \pm 0.220$ | $0.355 \pm 0.092$ |
| | $u_2$ (day<sup>-1</sup>) | $1.521 \pm 0.694$ | $3.839 \pm 0.576$ | $2.672 \pm 0.739$ |
| | $T_1$ (day) | $4.3 \pm 0.4$ | $4.3 \pm 0.3$ | $4.8 \pm 0.4$ |
| | $T_2$ (day) | $8.1 \pm 0.5$ | $8.2 \pm 0.6$ | $8.5 \pm 0.4$ |
|  | Ave. NMSE | 0.0005 | 0.0025 | 0.0009 |
|  | Max. NMSE | 0.0026 | 0.0091 | 0.0056 |
| Radiotherapy<br>at 8 Gy | $\lambda$ (day<sup>-1</sup>) | $0.490 \pm 0.041$ | $0.768 \pm 0.068$ | $0.465 \pm 0.046$ |
| | $u_1$ (day<sup>-1</sup>) | $0.997 \pm 0.110$ | $1.687 \pm 0.254$ | $0.787 \pm 0.134$ |
| | $u_2$ (day<sup>-1</sup>) | $2.147 \pm 0.627$ | $3.323 \pm 0.594$ | $2.815 \pm 0.707$ |
| | $T_1$ (day) | $3.8 \pm 0.3$ | $4.3 \pm 0.3$ | $4.5 \pm 0.3$ |
| | $T_2$ (day) | $7.9 \pm 0.5$ | $8.4 \pm 0.4$ | $8.5 \pm 0.4$ |
|  | Ave. NMSE | 0.0013 | 0.0028 | 0.0013 |
|  | Max. NMSE | 0.0055 | 0.0107 | 0.0089 |

**Figure 1.**
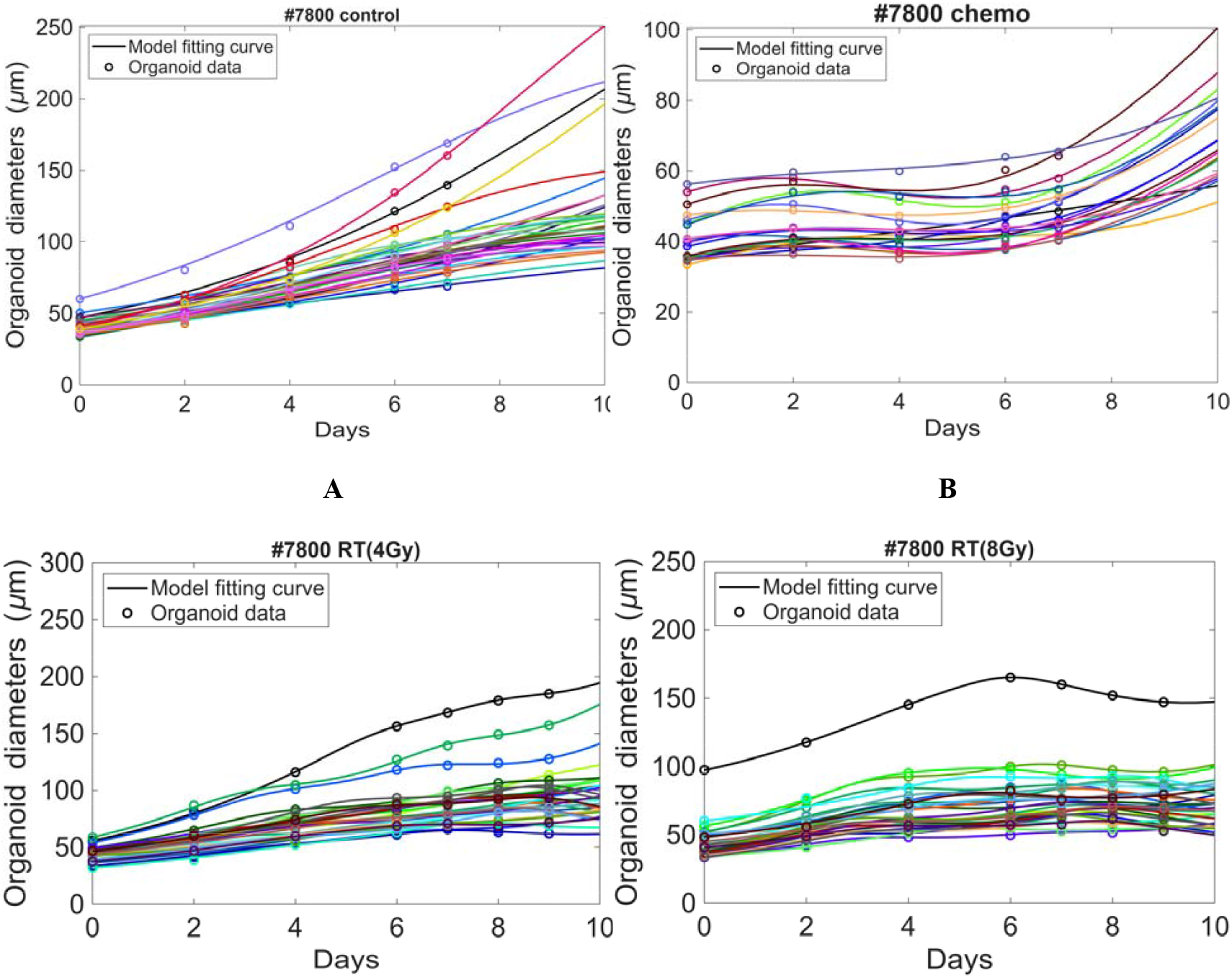
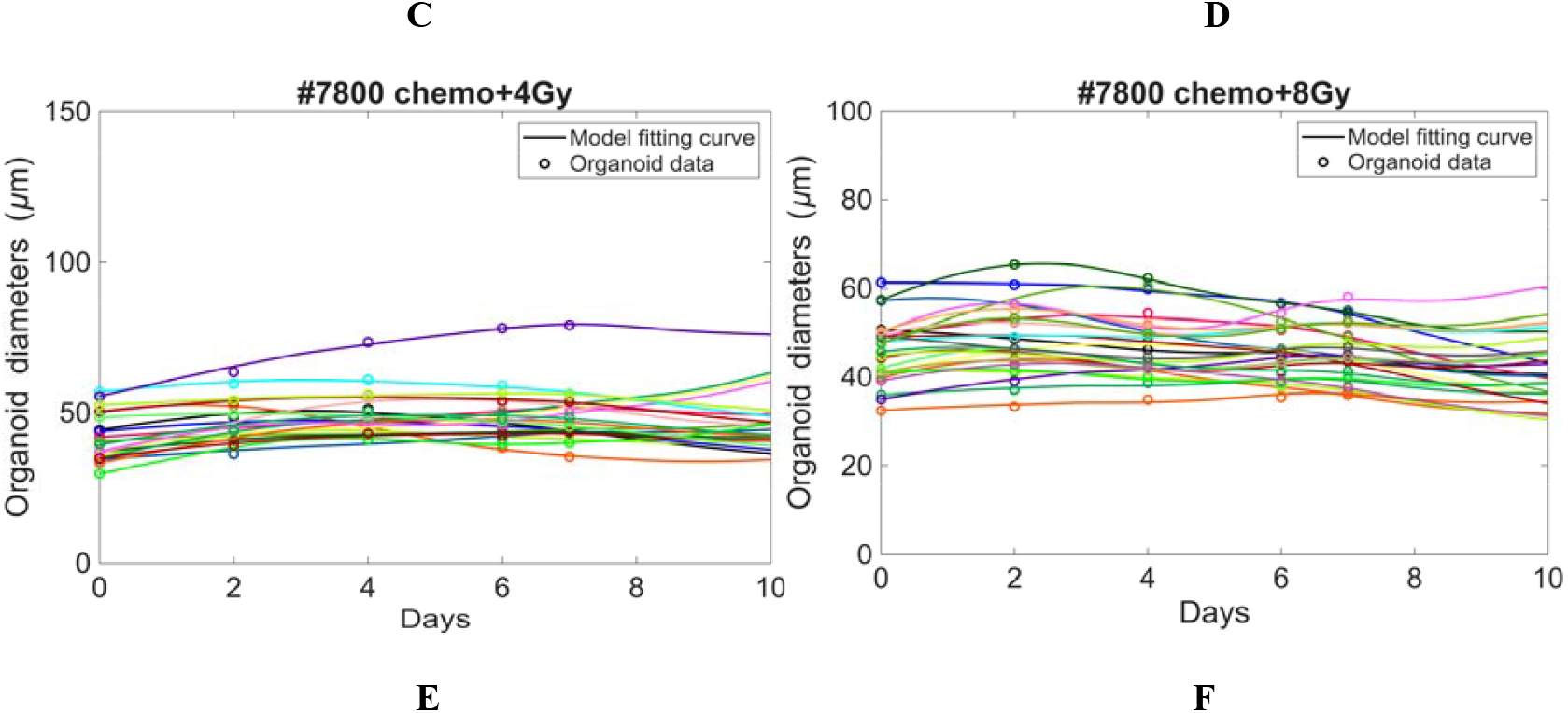
Model curve fittings of organoids #7800 in each group. Solid lines are fitted curves, and circles of the same color represent data from one organoid collected on corresponding days. There are 33, 20, 30, 30, 21, and 26 organoids’ size data collected in the control, chemotherapy, radiotherapy with 4 Gy, radiotherapy with 8 Gy, chemoradiotherapy with 4 Gy, and chemoradiotherapy with 8 Gy groups, respectively. The data points were taken up to 7 or 9 days for each group.

### 3.2 Chemotherapy group

We applied model (2) to the data of PDTOs in the chemotherapy group on days 0, 2, 4, 6, and 7 after a dose of drugs. Using the average growth rate *λ* and maximum carrying capacity *K* estimated from the control group, we fit the chemo-induced killing rate *C*(*t*), which is determined by the parameters *a,b*, and *T*_*c*_**. Figures 1(B), S2(B)**, and **S3(B)** show the fitting curves with the collected data for organoids #7800, #8510, and #11777, respectively. The fitting parameter values with 95% CIs and goodness of fit are all summarized in **Table 1 (Chemotherapy)**. These results indicate that the slowest-growing organoids #11777 exhibit the strongest drug resistance, with the smallest *a*-value (0.395) and a delayed peak killing effect occurring around day 5, compared with approximately day 4 for the other two samples. In contrast, the fastest-growing organoids #8510 are the most drug-sensitive (*a* = 0.906) but have a short killing-effect window (*b* = 0.162). Organoids #7800 display the most intense initial drug response, coupled with moderate sensitivity and a moderate-length killing-effect window. **Figure 2(A)** illustrates the time-dependent changes in killing rate C(t) for each organoid sample: the most sensitive organoids #8510 show a brief effect lasting about 10 days, the most resistant organoids #11777 sustain an effect for over 12 days, and #7800 fall between these extremes. These findings align with clinical practice, where each cycle of FOLFIRINOX chemotherapy typically spans approximately two weeks [22], and may reveal a possible relationship between the natural growth rate and drug resistance for these organoid samples; slower-growing organoids may exhibit stronger drug resistance but longer-lasting killing effect, primarily due to the development of resistance mechanisms and the heterogeneity of the pancreatic tumor itself [23].

**Figure 2.**
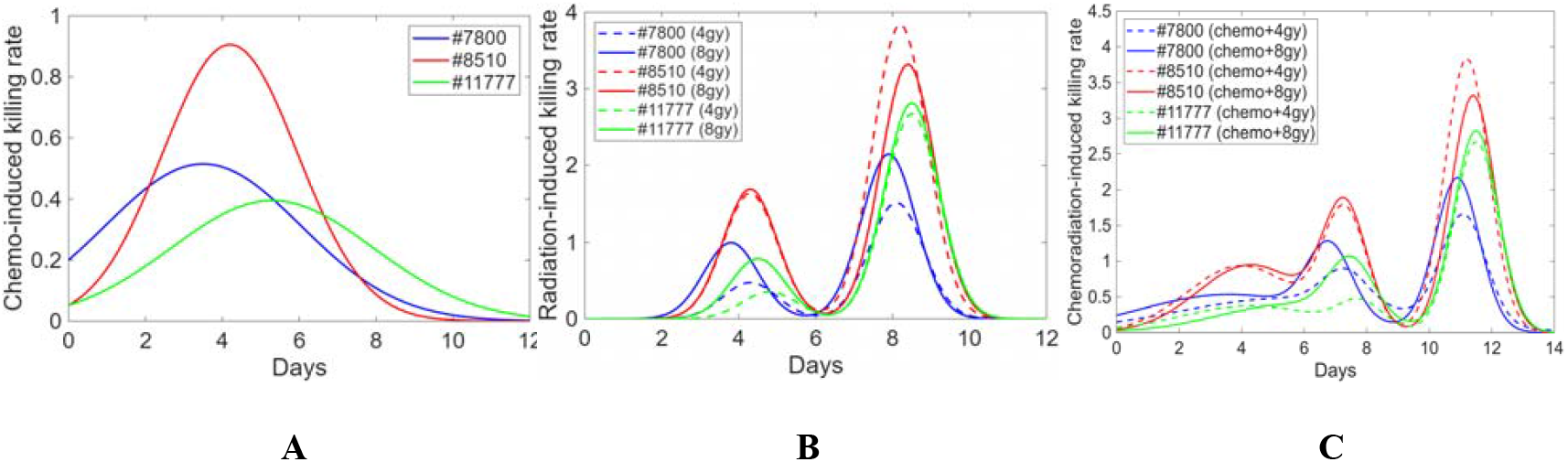
Treatment killing effect over time for #7800, #8510, and #11777 organoids. **A:** The chemo-induced killing effect occurs within days 0-12, with the response peak occurring around days 3-6. **B:** The radiation-induced killing effect (dashed lines for 4 Gy and solid lines for 8 Gy) occurs within days 2-11, with the early and secondary response mainly peaking around day 4 and day 8, respectively. **C:** The synergistic killing effect of chemoradiotherapy (dashed lines for 4 Gy and solid lines for 8 Gy) may last for the entire two weeks, with the early and secondary response primarily peaking around days 4-8 and days 11-12, respectively.

### 3.3 Radiotherapy group

For the radiotherapy group, we applied model (3) to organoid size measurements on days 0, 2, 4, 6, 7, 8, and 9 following radiation doses of 4 Gy and 8 Gy, respectively. In this group, the maximum carrying capacity *K* for each sample was fixed, and the removal rate was set to *μ*=0.3, within the typical range of 0 - 0.5 [24, 25]. The growth rates of active cells and radiation-induced killing rate R(t) were estimated directly from the data. The radiosensitivity parameters in the LQ model were set to *α* = 0.015 and *β* =*α*/9.5 [26] to estimate the surviving cell fraction. **Figures 1(C-D), S2(C-D)**, and **S3(C-D)** compare the fitted curves with the collected data of organoid samples at both radiation doses. **Table 1 (Radiotherapy)** presents the fitted growth rates of active cells and all parameters associated with radiation-induced killing rates. Across all organoid samples and both dose groups, the average early-response peak timing (*T*_1_) occurs around days 3.5-5, and the interval between *T*_1_ and the average secondary-response peak timing *T*_2_ was approximately 4 days, with the average secondary response strength consistently exceeding that of the early response (*u*_2_ > *u*_1_), possibly due to a combined effect of delayed apoptosis, cell cycle arrest, bystander effects, microenvironment modulation, etc. [27, 28]. For organoids #7800 and #11777, increasing the dose from 4 Gy to 8 Gy resulted in higher growth rates of active cells and earlier response timings which could be due to radiation-induced adaptive-response. In contrast, #8510 appeared less sensitive to dose escalation, showing comparable growth rates and peak timings at both doses, and even exhibiting a slightly higher killing rate at 4 Gy than at 8 Gy, suggestive of potential hyper-radiosensitivity (HRS) at low doses [29]. Nevertheless, organoids #8510 displayed the highest radiosensitivity, with *u*_1_ and *u*_2_ values significantly exceeding those of the other two organoid samples. Using the parameter values in **Table 1**, the overall radiation-induced killing rates for all three organoid samples are illustrated in **Figure 2(B)**. The early response lasted approximately 4 days, beginning at 2 - 3 days post-irradiation for 4 Gy and 1.5 - 2.5 days for 8 Gy. The secondary response persisted slightly longer, for about 4.5 days. Among these three organoid samples, #7800 was the most responsive to dose escalation: at 8 Gy, their peak responses occurred about 0.5 days earlier, and the magnitudes of both response waves increased markedly compared with 4 Gy. For #8510, early responses were similar for both doses, whereas the secondary response was slightly stronger at 4 Gy. Organoids #11777 exhibited the slowest onset, with the weakest early response but a relatively stronger secondary response, possibly caused by mitotic catastrophe, a form of cell death that occurs independently of typical apoptosis markers, potentially providing an escape route for apoptosis-resistant tumor cells like #11777 that depend on alternative pathways for controlling aberrant proliferation [30, 31]. This indicates that slower-growing organoids may exhibit a stronger resistance to radiation in the early phase, but not in the secondary phase. These findings highlight the complex intrinsic heterogeneity of PDTOs’ responses to radiation [20].

### 3.4 Chemoradiotherapy group

In the chemoradiotherapy group, the synergistic effects of chemotherapy and radiotherapy involve complex biological mechanisms, typically driven by multiple synergistic processes: radiotherapy can potentiate the efficacy of chemotherapy, while chemotherapy may enhance the radiosensitivity of tumor cells [32, 33]. To investigate the relative contributions of radiation-induced and chemo-induced killing in this synergistic interaction, we employed model (4) to separately fit the chemo-induced killing rate C(t) and radiation-induced killing rate R(t), while fixing the other component in each case.

In the first study, the parameters of the radiation-induced killing component R(t) were fixed in model (4) using the values determined in subsection 3.3, and the parameters of the chemo-induced killing component C(t) were estimated by fitting to the chemoradiotherapy data. **Figures 1(E-F), S2(E-F)**, and **S3(E-F)** compare the fitted curves with the collected data of organoid samples, yielding low average NMSE values (all below 0.0030) as shown in **Table 2**, indicating a reasonably good fit. **Figure 2(C)** presents the total synergistic killing rate of chemoradiotherapy for each organoid sample. It shows that combined therapy yields a longer and more intense killing window than either modality alone. In particular, the response during the first week was substantially improved across all organoid types compared to radiotherapy or chemotherapy alone, while the secondary response resembled that observed with radiotherapy only, suggesting that the synergistic effect may primarily occur during the first week post radiation.

**Table 2.** Fitting results for the chemo-induced killing rate C(t) and radiation-induced killing rate R(t) in the chemoradiotherapy group, with one fixed and the other fitted.

|  | PDTOs ID | #7800 | #8510 | #11777 |
| --- | --- | --- | --- | --- |
| Fitting $C(t)$ with<br>fixed $R(t)$ at 4 Gy | $a$ (day<sup>-1</sup>) | $0.482 \pm 0.042$ | $0.939 \pm 0.115$ | $0.374 \pm 0.032$ |
| | $b$ | $0.039 \pm 0.019$ | $0.167 \pm 0.054$ | $0.096 \pm 0.048$ |
| | $T_c$ (day) | $5.5 \pm 0.7$ | $4.0 \pm 0.3$ | $4.3 \pm 0.6$ |
|  | Ave. NMSE | 0.0017 | 0.0027 | 0.0024 |
|  | Max. NMSE | 0.0060 | 0.0106 | 0.0082 |
| Fitting $R(t)$ with<br>fixed $C(t)$ at 4 Gy | Ave. NMSE | 0.0837 | 0.2185 | 0.0706 |
|  | Max. NMSE | 0.1438 | 0.7684 | 0.2439 |
| Fitting C(t) with<br>fixed R(t) at 8 Gy | $a$ (day<sup>-1</sup>) | $0.536 \pm 0.032$ | $0.950 \pm 0.092$ | $0.396 \pm 0.032$ |
| | $b$ | $0.061 \pm 0.020$ | $0.173 \pm 0.064$ | $0.093 \pm 0.049$ |
| | $T_c$ (day) | $3.6 \pm 0.6$ | $4.3 \pm 0.5$ | $5.6 \pm 0.6$ |
|  | Ave. NMSE | 0.0009 | 0.0029 | 0.0020 |
|  | Max. NMSE | 0.0032 | 0.0105 | 0.0088 |
| Fitting R(t) with<br>fixed C(t) at 8 Gy | Ave. NMSE | 0.0330 | 0.1930 | 0.0437 |
|  | Max. NMSE | 0.1194 | 0.6276 | 0.1726 |

In the second study, the parameters of the chemo-induced killing rate C(t) were fixed in model (4) using values determined in subsection 3.2, and the parameters of the radiation-induced killing rate R(t) were estimated by fitting to the same dataset from the chemoradiotherapy group above. The resulting fits were much inferior to those obtained in the first study, as shown by the substantially higher average and maximum NMSE values for each organoid sample in **Table 2**, where the parameter values were thus not provided in this case.

These two studies suggest that the radiation-induced killing effect remains relatively stable, while the chemo-induced killing effect changes when comparing monotherapies with combined chemoradiotherapy. This indicates that the influence of radiotherapy on chemotherapy, such as modulating tumor microenvironment and DNA repair pathways to make cells more vulnerable to drugs, is more pronounced than that of chemotherapy on radiotherapy, such as interfering with DNA repair mechanisms to cause cell cycle arrest in radiation-sensitive phases [34]. In other words, the radiation-induced effect may be the dominant driver of the synergistic interaction observed across the three organoid samples, which also exhibit heterogeneity in the change of chemo-induced effect. Specifically, **Table 2** demonstrates that an 8 Gy dose universally enhanced chemotherapy efficacy compared to 4 Gy (characterized by higher *a*). Comparison between the chemotherapy fitting results for chemotherapy alone in **Table 1** and chemoradiotherapy in **Table 2** reveals that radiation therapy prolonged the chemo therapeutic window in #7800 (lower *b*), shortened the therapeutic window in #1177 (higher *b*), and increased the chemoradiotherapy in **Table 2** reveals that radiation therapy prolonged the chemo therapeutic window in chemosensitivity with 8 Gy radiation dose in #7800 and #8510 (higher *a*).

### 3.5 Comparison of treatment efficacy

**Figures 2(A-C)** show that the organoids displayed an earlier initial response to chemotherapy than to radiotherapy; however, two markedly stronger responses, especially the second one, to radiotherapy emerged during the following two weeks. The combination of chemotherapy and radiotherapy extended the overall treatment response across nearly two weeks post-treatment, characterized by a longer and stronger early phase caused by synergistic interactions, followed by a pronounced secondary phase driven by radiation. Moreover, organoids exhibiting greater chemosensitivity, such as #8510, also seem to display a stronger response to radiation. **Table 3** presents the overall treatment response (TR) on day 7 and the further TR of radiotherapy on day 9 based on the observed data. It is calculated as the logarithm of the ratio between the post-treatment organoid size from the data and the non-treated organoid size estimated by model (1) at the time of efficacy assessment, showing a continuous and dimensionless metric that captures the organoid response to both cytotoxic and cytostatic therapeutic effects [35]. A larger TR value indicates more effective therapy. It shows that chemotherapy and chemoradiotherapy have significantly larger values of TR than those of radiotherapy on day 7, and the TR on day 9 is much improved by at least 50% or so from day 7 for radiotherapy. These findings further support that chemotherapy and chemoradiotherapy trigger a stronger killing effect than radiotherapy in the first week, but radiotherapy may have a stronger second killing effect in the following week. Using the fitted parameters in each group, we simulated predicted growth curves for all three organoid samples, initiated from a diameter of 100 μm under different therapeutic regimens, shown in **Figure S4**, which illustrates the heterogeneity in therapeutic responses among organoid samples for 12 days post-treatment. In terms of therapeutic effects, chemotherapy induces a stronger response than radiotherapy during the first week (day 7); however, its efficacy progressively diminishes thereafter, with all organoid samples exhibiting marked regrowth. By contrast, radiotherapy sustains organoid growth inhibition into the following week due to the secondary killing effect and hence produces a favorable treatment effect (day 12) than chemotherapy. Notably, chemoradiotherapy integrates the benefits of both modalities, producing slightly greater efficacy than chemotherapy in the first week and significantly superior efficacy compared with radiotherapy in the second week. Specifically, organoids #7800 exhibit a moderate growth rate and a relatively small carrying capacity, and demonstrate sensitivity to both chemotherapy and radiotherapy (including dose variations); organoids #8510 grow rapidly, possess a larger carrying capacity, are highly sensitive to both modalities, and display HRS at a low dose; organoids #11777 grow slowly but have a comparatively large carrying capacity, and show partial resistance to both chemotherapy and radiotherapy during the early treatment phase, but exhibit increased sensitivity to radiation in the secondary response phase. Based on these findings, chemoradiotherapy at 8 Gy appears to provide the most effective tumor control for #7800 and some improvement over 4 Gy for #11777, but 4 Gy of chemoradiotherapy may be a sufficiently effective strategy for #8510.

**Table 3.** The treatment responses with 95% CIs on day 7 and day 9 for all three organoid samples.

| PDTOs ID |  | Day 7 | Day 9 |
| --- | --- | --- | --- |
| #7800 | Chemotherapy | 2.71 ± 0.12 |  |
|  | Radiotherapy at 4 Gy | 1.24 ± 0.15 | 2.15 ± 0.15 |
|  | Radiotherapy at 8 Gy | 1.62 ± 0.12 | 2.52 ± 0.12 |
|  | Chemoradiotherapy at 4Gy | 2.76 ± 0.25 |  |
|  | Chemoradiotherapy at 8Gy | 3.23 ± 0.13 |  |
| #8510 | Chemotherapy | 3.79 ± 0.21 |  |
|  | Radiotherapy at 4 Gy | 2.01 ± 0.19 | 3.00 ± 0.24 |
|  | Radiotherapy at 8 Gy | 1.96 ± 0.18 | 3.01 ± 0.23 |
|  | Chemoradiotherapy at 4Gy | 4.03 ± 0.18 |  |
|  | Chemoradiotherapy at 8Gy | 3.97 ± 0.16 |  |
| #11777 | Chemotherapy | 1.98 ± 0.15 |  |
|  | Radiotherapy at 4 Gy | 0.69 ± 0.15 | 1.54 ± 0.15 |
|  | Radiotherapy at 8 Gy | 1.06 ± 0.15 | 1.92 ± 0.15 |
|  | Chemoradiotherapy at 4Gy | 1.89 ± 0.17 |  |
|  | Chemoradiotherapy at 8Gy | 2.01 ± 0.16 |  |

## 4. Discussion

Patient-derived tumor organoids offer substantial advantages for personalized cancer treatment, as they closely recapitulate the biological and molecular characteristics of the original patient tumors. Their use enables more accurate and individualized treatment testing and selection, thereby enhancing the flexibility and reliability of preclinical evaluations. This platform holds significant potential for optimizing and refining treatment strategies, ultimately improving clinical outcomes [36]. Effective cancer therapy often depends on a combination of modalities, such as radiotherapy and/or chemotherapy, where dose, fractionation, sequencing, and other parameters can critically influence treatment efficacy [37, 38]. However, evaluating additional therapeutic strategies in clinical trials is both time-consuming and costly. Mathematical modeling offers a powerful and efficient approach to exploring a broader range of clinical options using limited experimental or clinical data, thereby informing and optimizing the design of personalized treatment plans. The results obtained through modelling have shown that tumor organoids #11777 displayed resistance to chemotherapy drugs and radiation. We also compared this data with the limited medical history of the patient #11777, who was treatment naive at the time of disease diagnosis and could have developed intrinsic resistance to chemotherapy drugs and radiation due to a mutation in DNA repair genes [39, 40].

We developed mathematical models based on the logistic growth framework to characterize the killing effects of chemotherapy, radiotherapy, and chemoradiotherapy in distinct PDTOs. The presented models accurately captured the growth dynamics of each organoid sample under different therapeutic regimens using a limited set of parameters. Notably, they provided, to the best of our knowledge, the first quantitative assessment of the individual contribution of each therapy and the synergistic process of chemoradiotherapy, while previous studies have primarily focused on overall response assessments of PDTOs to various therapies [11, 16, 20], and our modeling approach enables rapid and mechanistic insights into post-treatment growth dynamics. Furthermore, it highlights interpatient variability in treatment responses, thereby supporting the design and optimization of clinical trials and personalized tumor treatment planning in precision medicine.

Our models focus on the growth dynamics of PDTOs after treatment and do not explicitly account for molecular mechanisms such as DNA repair, cell-cycle effects, or drug resistance, which are key determinants of therapeutic outcomes [41, 42]. Nevertheless, they effectively capture organoids’ growth dynamics over time and quantify responses to chemotherapy and radiotherapy through the chemo-induced and radiation-induced killing effects, C(t) and R(t), as well as their synergistic interaction. The results underscore pronounced heterogeneity in treatment responses across different PDTOs. Despite these differences, chemoradiotherapy consistently enhances tumor control by inducing earlier onset and stronger killing effects during the first week compared with either modality alone. These findings may inform therapy optimization in precision medicine. Noticeably, future work could be extended to larger and more diverse cohorts by integrating molecular and spatial mechanisms to reflect genetic and phenotypic heterogeneity and evaluate long-term responses to strengthen translational relevance and support personalized therapy design.

## 5. Conclusion

We proposed mathematical frameworks that separately characterize the cytotoxic effects of chemotherapy, radiotherapy, and their potential synergy in combined chemoradiotherapy. Each model demonstrated a strong fit to the respective treatment group data. Through model fitting and simulation, we quantitatively captured key features of treatment-induced cell killing, including killing strength, onset timing, and duration, revealing that the effect of chemotherapy is predominantly concentrated in the first week, whereas that of radiotherapy is more pronounced in the following week. Additionally, we used a mathematical approach to elucidate the synergistic interactions between radiation and chemotherapeutic agents, demonstrating the dominant impact of radiation therapy on chemotherapy in these interactions. Our results also revealed substantial heterogeneity in the growth dynamics and therapy responses of PDTOs. These findings underscore the potential of PDTOs as a preclinical platform for guiding treatment selection and informing clinical trial design to optimize the efficacy of chemotherapy, radiotherapy, and combined chemoradiotherapy regimens in precision medicine.

## Supplementary Material

**Figure S1.**
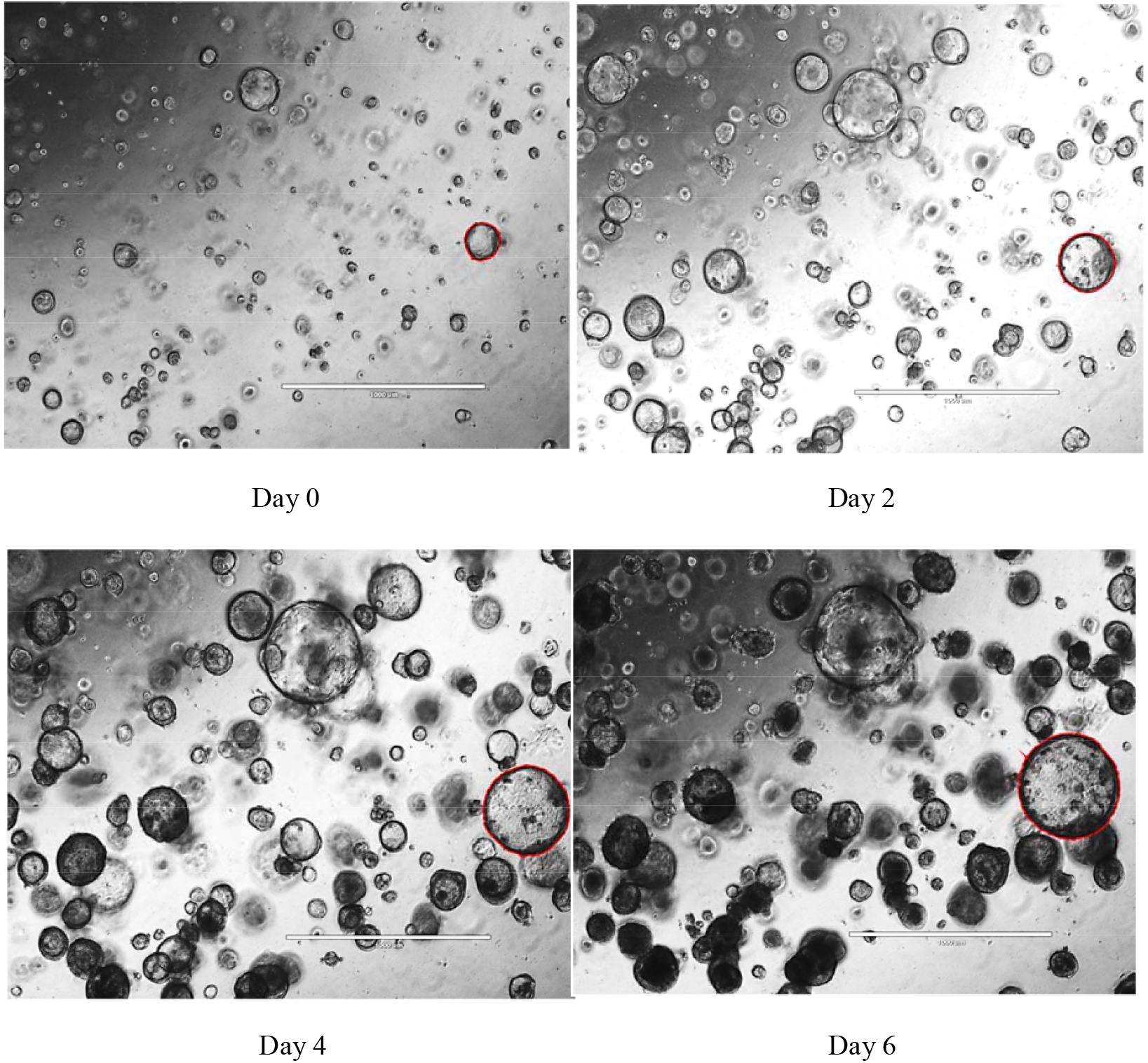
Brightfield image samples of organoids #8510 on days 0, 2, 4, and 6 after radiation of 4 Gy. The typical organoid size was determined by area *S* enclosed by the red circle, that is, the organoid diameter is calculated by 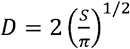. Scale bar = 1000 μm.

**Figure S2.**
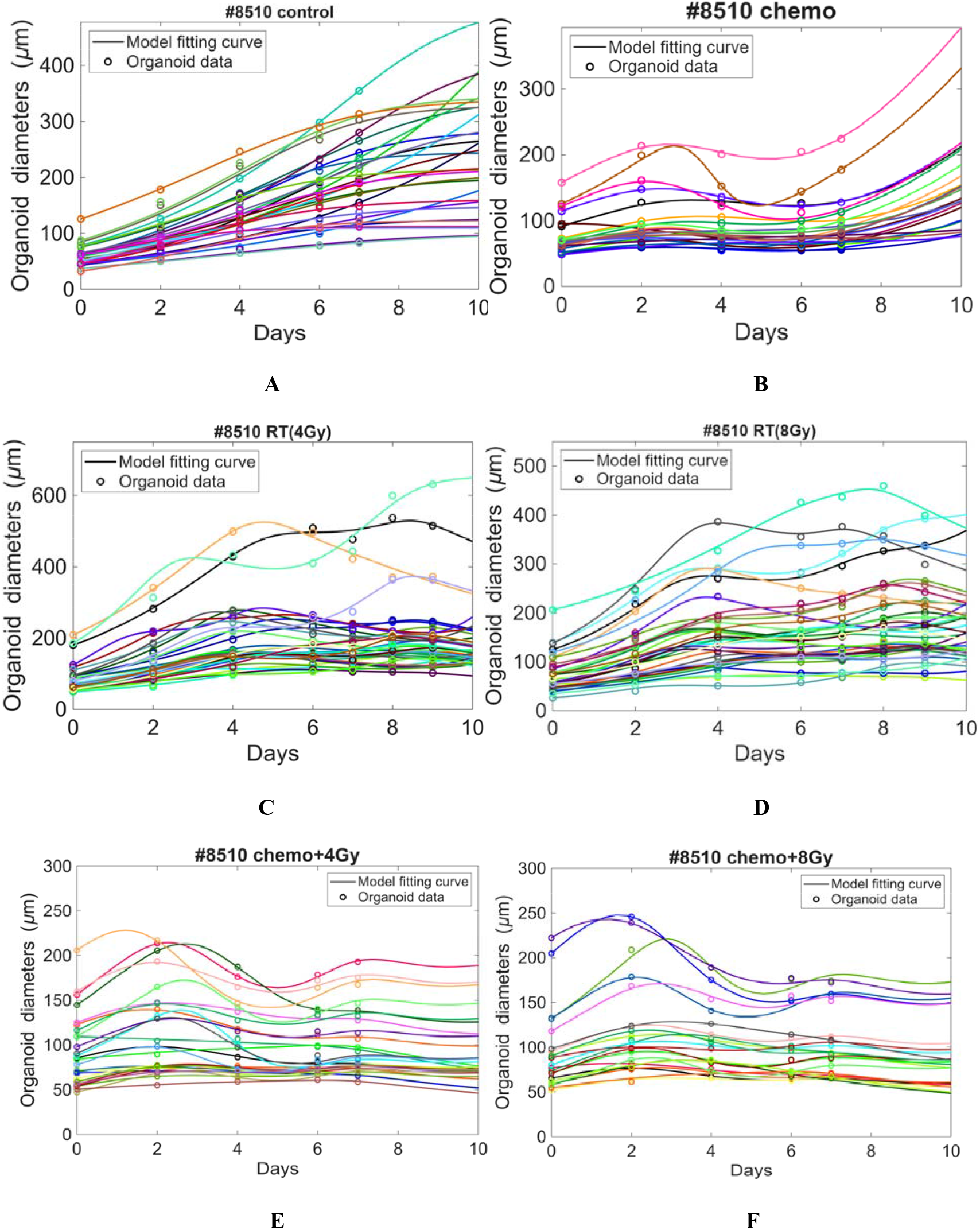
Model curve fittings of organoids #8510 in each group. Solid lines are fitted curves, and circles of the same color represent data from one organoid collected on corresponding days. There are 30, 24, 37, 37, 28, and 22 organoid size data collected in the control, chemotherapy, radiotherapy with 4 Gy, radiotherapy with 8 Gy, chemoradiotherapy with 4 Gy, and chemoradiotherapy with 8 Gy groups, respectively. The data points were taken up to 7 or 9 days for each group.

**Figure S3.**
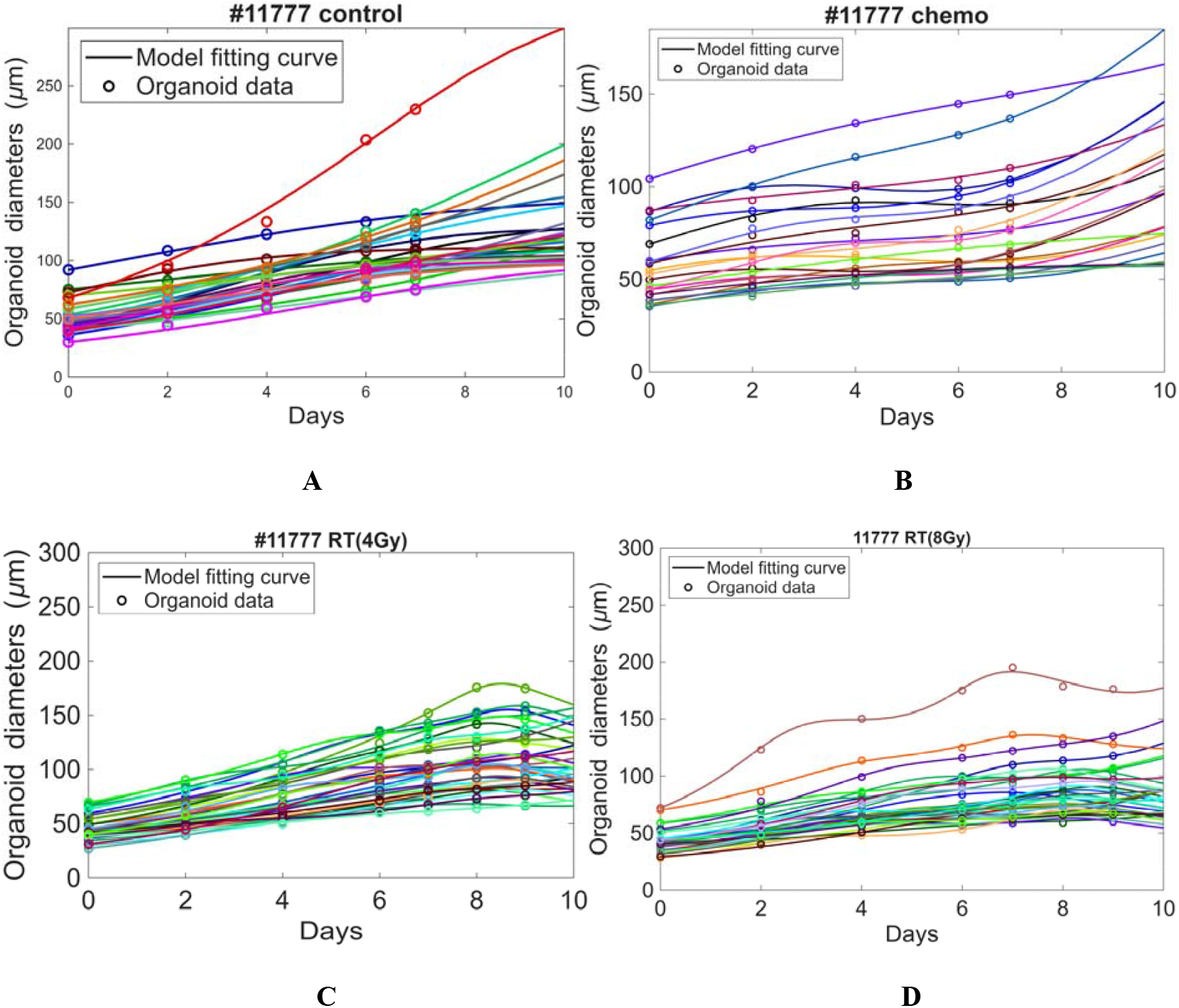
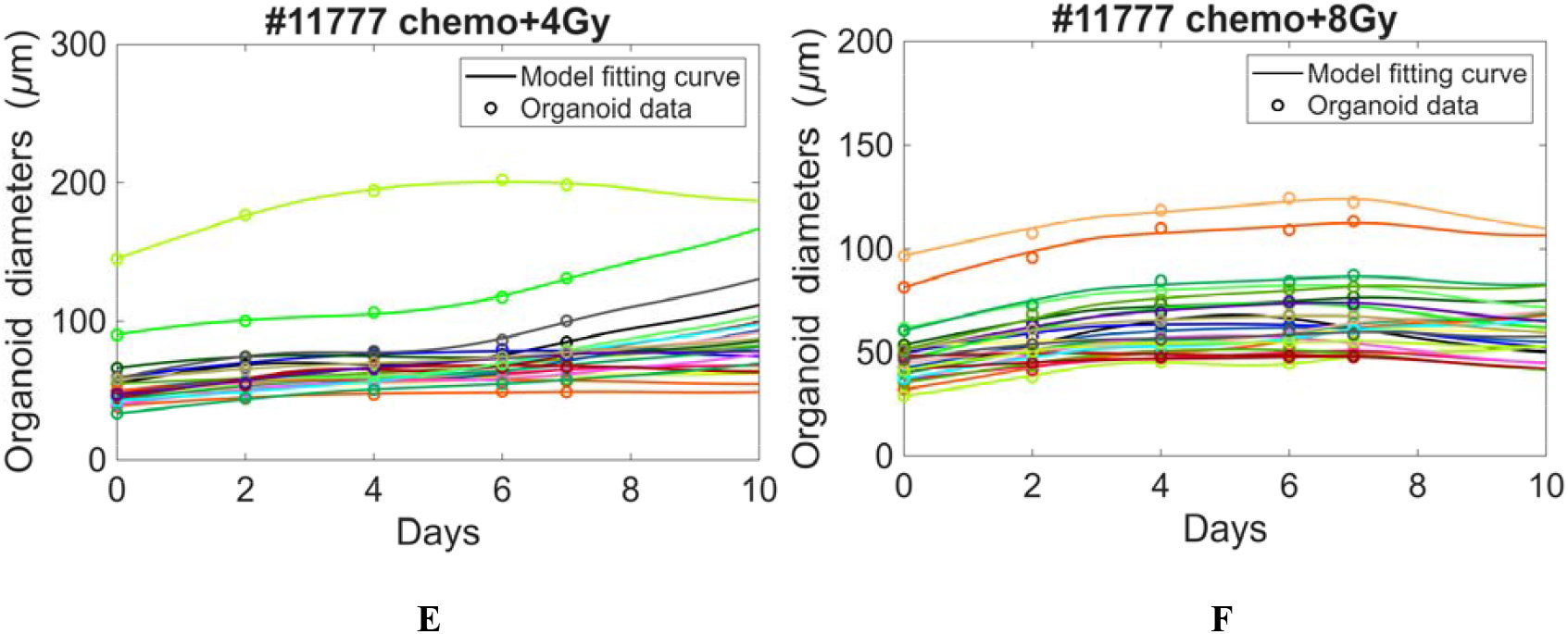
Model curve fittings of organoids #11777 in each group. Solid lines are fitted curves, and circles of the same color represent data from one organoid collected on corresponding days. There are 30, 34, 35, 22, 23, and 25 organoid size data collected in the control, chemotherapy, radiotherapy with 4 Gy, radiotherapy with 8 Gy, chemoradiotherapy with 4 Gy, and chemoradiotherapy with 8 Gy groups, respectively. The data points were taken up to 7 or 9 days for each group.

**Figure S4.**
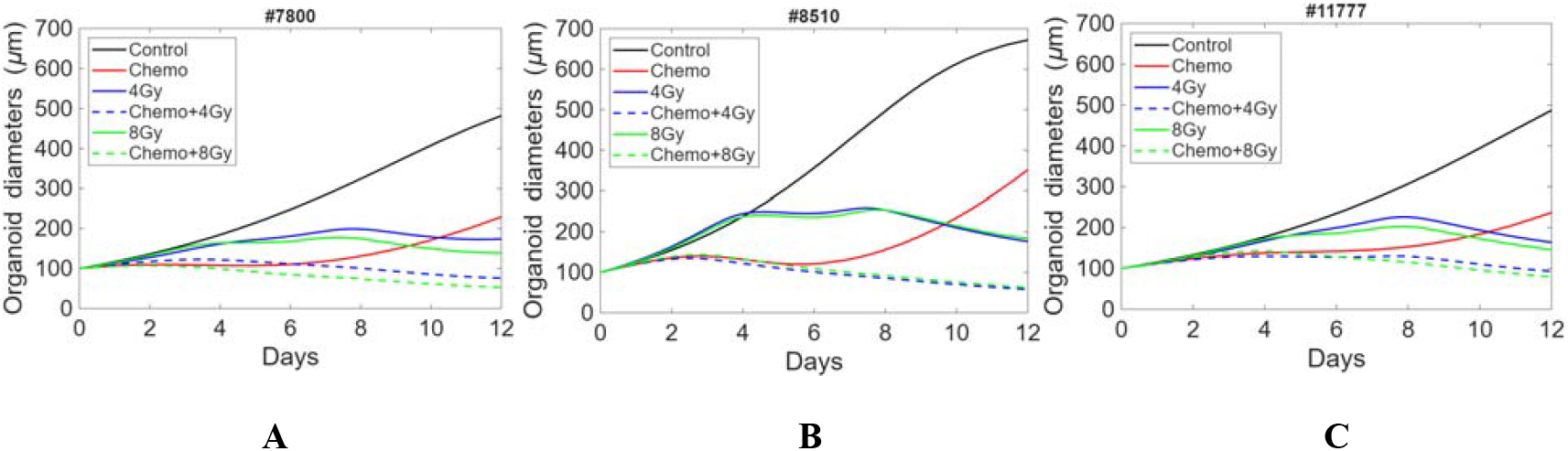
Simulated organoid diameters in each sample starting at 100 *μm* change over time for up to 12 days in the scenarios of control, chemotherapy, radiotherapy with 4 Gy and 8 Gy, and chemoradiotherapy with 4 Gy and 8 Gy.

**Table S1.** The predicted overall effect of treatment responses (TR) on day 12.

| PDTOs ID | #7800 | #8510 | #11777 |
| --- | --- | --- | --- |
| Chemotherapy | 2.23 | 1.94 | 2.16 |
| Radiotherapy at 4 Gy | 3.06 | 4.01 | 3.27 |
| Radiotherapy at 8 Gy | 3.73 | 3.89 | 3.61 |
| Chemoradiotherapy at 4 Gy | 5.54 | 7.36 | 4.97 |
| Chemoradiotherapy at 8 Gy | 6.63 | 7.14 | 5.44 |

## References

[1] Ma X, Wang Q, Li G, et al. Cancer organoids: A platform in basic and translational research. Genes Dis. 2023, 11(2):614–632.

[2] Yang H, Wang Y, Wang P, et al. Tumor organoids for cancer research and personalized medicine. Cancer biology & medicine 2022, 19(3) 319-332.

[3] Qu S, Xu R, Yi G, et al. Patient-derived organoids in human cancer: a platform for fundamental research and precision medicine. Mol Biomed. 2024, 5(1):6.

[4] Foo MA, You M, Chan SL, et al. Clinical translation of patient-derived tumor organoids-bottlenecks and strategies. Biomarker Research 10, 10 (2022).

[5] Wensink GE, Elias SG, Mullenders J, et al. Patient-derived organoids as a predictive biomarker for treatment response in cancer patients. npj Precision Oncology 5, 30 (2021).

[6] Wang J, Chen C, Wang L, et al. Patient-Derived Tumor Organoids: New Progress and Opportunities to Facilitate Precision Cancer Immunotherapy. Front Oncol. 2022, 12:872531.

[7] Pasch CA, Favreau PF, Yueh AE, et al. Patient-Derived Cancer Organoid Cultures to Predict Sensitivity to Chemotherapy and Radiation. Clinical Cancer Research. 25(17), 2019.

[8] Xu H, Jiao D, Liu A, and Wu K. Tumor organoids: applications in cancer modeling and potentials in precision medicine. Journal of Hematology & Oncology (2022) 15:58.

[9] Wang Y and Jeon H. (2022) 3D cell cultures toward quantitative high-throughput drug screening. Trends in Pharmacological Sciences, 43(7):569–581.

[10] van de Wetering M, Francies HE, Francis JM, et al. (2015) Prospective Derivation of a Living Organoid Biobank of Colorectal Cancer Patients. Cell, 161(4):933–945.

[11] Shukla HD, Dukic T, Roy S, et al. (2023) Pancreatic cancer derived 3D organoids as a clinical tool to evaluate the treatment response. Front. Oncol. 12:1072774.

[12] Roy S, Dukic T, Keepers Z, et al. SOX2 and OCT4 mediate radiation and drug resistance in pancreatic tumor organoids. Cell Death Discovery (2024)10:106.

[13] Ganesh K, Wu C, O’Rourke KP, et al. A rectal cancer organoid platform to study individual responses to chemoradiation. Nature Medicine 25, 1607–16114, 2019.

[14] Tiriac H, Belleau P, Engle DD, et al. Organoid Profiling Identifies Common Responders to Chemotherapy in Pancreatic Cancer. Cancer Discovery (2018), 8(9):1112–1129.

[15] Drost J, Clevers H. Organoids in cancer research. Nature Reviews Cancer 18, 407–418, 2018.

[16] Yao Y, Xu X, Yang L, et al. Patient-Derived Organoids Predict Chemoradiation Responses of Locally Advanced Rectal Cancer. Clinical and Translational Report 26, 1:17–26, 2020.

[17] Gunnarsson EB, Kim S, Choi B, et al. (2024) Understanding patient-derived tumor organoid growth through an integrated imaging and mathematical modeling framework. PLOS computational biology 20(8): e1012256.

[18] Kim JS, Park CH, Kim E, et al. (2025) Establishing 3D organoid models from patient-derived conditionally reprogrammed cells to bridge preclinical and clinical insights in pancreatic cancer. Mol Cancer 24, 162.

[19] Nicosia L, Alongi F, Andreani S, et al. Combinatorial effect of magnetic field and radiotherapy in PDAC organoids: a pilot study. Biomedicines 2020;8(12):609.

[20] Iris WJM van Goor, Leon Raymakers, Daan SH Andel et al. Radiation response assessment of organoids derived from patients with pancreatic cancer. Clinical and Translational Radiation Oncology 48 (2024) 100829.

[21] Walter N, Micoulet A, Seufferlein T, Spatz JP. Direct assessment of living cell mechanical responses during deformation inside microchannel restrictions. Biointerphases 6, 117–125 (2011).

[22] Conroy T, Desseigne F, Ychou M, et al. FOLFIRINOX versus gemcitabine for metastatic pancreatic cancer. N Engl J Med. 2011, 364(19):1817–25.

[23] Le Compte M, De La Hoz EC, Peeters S. et al. Single-organoid analysis reveals clinically relevant treatment-resistant and invasive subclones in pancreatic cancer. npj Precis. Onc. 7, 128 (2023).

[24] Watanabe Y, Dahlman EL, Leder KZ, and Hui SK. A mathematical model of tumor growth and its response to single irradiation. Theoretical Biology and Medical Modelling, (2016) 13:6.

[25] Yang C, Li J, Li W et al. A predictive model studying the impact of timing and order for the combination-regime of spatially fractionated radiation therapy and stereotactic body radiation therapy. Phys. Med. Biol. 70 (2025) 145016.

[26] Prior PW, Chen X, Hall WA, et al. (2018) Estimation of the Alpha-beta Ratio for Chemoradiation of Locally Advanced Pancreatic Cancer. International Journal of Radiation Oncology, Biology, Physics, Volume 102, Issue 3, S97.

[27] Kim JH, Brown SL, Gordon MN. Radiation-induced senescence: therapeutic opportunities. Radiation Oncology 18, 10 (2023).

[28] Daguenet E, Louati S, Wozny, AS. et al. Radiation-induced bystander and abscopal effects: important lessons from preclinical models. Br J Cancer 123, 339–348 (2020).

[29] Joiner MC, Marples B, Lambin P, et al. (2001) Low-dose hypersensitivity: current status and possible mechanisms. International Journal of Radiation Oncology, Biology, Physics, Volume 49, Issue 2, 319–389.

[30] Yoshino Y and Ishioka C. Inhibition of glycogen synthase kinase-3 beta induces apoptosis and mitotic catastrophe by disrupting centrosome regulation in cancer cells. Scientific Reports 5, 13249 (2015).

[31] Sia J, Szmyd R, Hau E, et al. Molecular Mechanisms of Radiation-Induced Cancer Cell Death: A Primer. Front. Cell Dev. Biol, 8:41 (2020).

[32] Choy H and Kim DW. (2003) Chemotherapy and irradiation interaction. Seminars in Oncology. Volume 30, S9, 3–10.

[33] Rallis KS, Yau THL, and Sideris M. (2021) Chemoradiotherapy in cancer treatment: rationale and clinical applications. Anticancer Research. 2021, 41(1), 1–7.

[34] Wang Y, Li Y, Sheng Z, et al. (2022) Advances of Patient-Derived Organoids in Personalized Radiotherapy. Front. Oncol. 12:888416.

[35] Mehrara E, Forssell-Aronsson E, Bernhardt P. Objective assessment of tumor response to therapy based on tumor growth kinetics. Br J Cancer 105, 682–686 (2011).

[36] Thorel L, Perréard M, and Florent R, et al. (2024) Patient-derived tumor organoids: a new avenue for preclinical research and precision medicine in oncology. Experimental & Molecular Medicine 56, 1531–1551.

[37] Lazzari G, Montagna A, D’Andrea B, et al. Breast Cancer Adjuvant Radiotherapy and Chemotherapy Sequencing: Sequential, Concomitant, or What Else? A Comprehensive Review of the Adjuvant Combinations Journey. J Clin Med. 2024, 13(20):6251.

[38] Yang YC and Chiang CS. Challenges of Using High-Dose Fractionation Radiotherapy in Combination Therapy. Front Oncol. 2016, 6:165.

[39] Keepers ZL, Roy S, William R, et al. Patient-derived pancreatic tumor organoids as a tool to evaluate cancer stem cell populations and their role in therapeutic resistance. Cancer Res 2024;84(17 Suppl_2):Abstract nr B086.

[40] Kawashima A, Takayama H, Tsujimura A. A Review of ERCC1 Gene in Bladder Cancer: Implications for Carcinogenesis and Resistance to Chemoradiotherapy. Adv Urol. 2012;2012:812398.

[41] Helleday T, Petermann E, Lundin C, et al. DNA repair pathways as targets for cancer therapy, Nat Rev Cancer 8, 193–204 (2008).

[42] Holohan C, Van Schaeybroeck S, Longley D, et al. Cancer drug resistance: an evolving paradigm. Nat Rev Cancer 13, 714–726 (2013).

